# Hypodopaminergic state of the nigrostriatal pathway drives compulsive alcohol use

**DOI:** 10.1101/2021.12.24.474124

**Authors:** Raphaël Goutaudier, David Mallet, Magali Bartolomucci, Denis Guicherd, Carole Carcenac, Frédérique Vossier, Thibault Dufourd, Sabrina Boulet, Colin Deransart, Benoit Chovelon, Sebastien Carnicella

## Abstract

The neurobiological mechanisms underlying compulsive alcohol use, a cardinal feature of alcohol use disorder, remain elusive. The key modulator of motivational processes dopamine (DA) is suspected to play an important role in this pathology, but its exact implication remains to be determined. Here, we found that rats expressing compulsive alcohol-related behavior, operationalized as punishment-resistant self-administration, showed a decrease in DA levels restricted to the dorsolateral territories of the striatum, the main output structure of the nigrostriatal DA pathway. We then causally demonstrated that a chemogenetic-induced selective hypodopaminergia of this pathway results in compulsive alcohol self-administration in rats otherwise resilient, accompanied by the emergence of alcohol withdrawal-like motivational impairments. These results demonstrate a major implication of tonic nigrostriatal hypodopaminergic state in alcohol addiction and provide new insights into our understanding of the neurobiological mechanisms underlying compulsive alcohol use.

## Introduction

More than 300 million people suffer from alcohol use disorder (AUD) worldwide (1), a pathology that encompasses a broad spectrum of health, social and economic problems with various degrees of severity (2). The most severe form is frequently referred as alcohol addiction, a chronic relapsing disease occurring only in a subset of vulnerable users (3). It is notably characterized by compulsive alcohol use, that is the continuation of seeking and drinking alcohol despite having significant negative consequences, and the presence of a negative motivational and affective state in absence of alcohol (2,4). While neuroscientists have identified a plethora of actors in AUD, treatments are limited mainly due to the poor understanding of the psychobiological mechanisms underlying the shift from controlled to compulsive alcohol seeking and taking (3). At the neural systems level, dopamine (DA) is considered as a prominent actor in the pathophysiology of addiction mainly through its role in incentive motivation and reinforcement (4). However, forty years of research have hitherto failed to provide a clear idea of its exact contribution (5). While alcohol consumption transiently increases DA levels in the ventral striatum, especially in the nucleus accumbens (NAc), the main output of the mesolimbic system (6), extended consumption instead leads to an overall and prolonged decrease of DA levels upon abstinence in this structure (7,8). Over the past decade, tonic hypodopaminergic states of the mesolimbic pathway have been causally linked to the emergence of both acute drug withdrawal symptoms and excessive alcohol seeking and taking (9,10). However, implication of these DA hypofunctions in protracted abstinence and in the compulsive dimension of AUD remains controversial (11,12).

Beyond the mesolimbic DA pathway, a growing body of research had recently demonstrated the strong implication in motivated and affective behaviors of the neighboring nigrostriatal DA pathway, originally restricted to motor functions (13,14). Interestingly, clinical studies in abstinent individuals with AUD not only have shown an altered DA function in the ventral striatum, but also within the dorsal striatum (8), the major output of the nigrostriatal DA pathway, that is assumed to play a prominent role in compulsive drug use (3,15–17). More specifically in rodents, compulsive alcohol seeking has been found to depend on the anterior part of the dorsolateral striatum (aDLS) and DA signaling in this structure (18). In striking contrast with the mesolimbic pathway however, study of potential tonic hypodopaminergic states of the nigrostriatal pathway in AUD has been neglected so far (16). We therefore hypothesized here that the existence of a nigrostriatal hypodopaminergia would contribute to the development of compulsive alcohol use and of a negative affective state.

## Materials and Methods

### Animals

Experiments were performed on *Wild-Type (WT)* and *TH::Cre* Long-Evans males rats. They were housed in a 12h/12h reverse light cycle, with food and water *ad libitum*. At the beginning of the experiments, rats were 7 to 8 weeks old. All experimental protocols complied with the European Union 2010 Animal Welfare Act and the new French directive 2010/63, and were approved by the French national ethics committee no. 004.

### Experimental design

#### Experiment 1

21 *WT* rats were trained during 26 sessions in an intermittent access to 20% alcohol two-bottle choice drinking procedure (IA 20%-EtOH 2BC). Operant ethanol self-administration was then performed during 38 sessions before the beginning of the footshock-punished ethanol self-administration. At this step, half of the rats performed 6 punished sessions followed by 6 unpunished sessions, while tests were carried out in opposite orders for the other half, thereby ensuring that any decrease in performance was the result of footshocks and not an effect of the time or environmental conditions. Additionally, 8 *WT* rats were kept under identical conditions but never experienced alcohol. Rats were euthanized after a period of abstinence (one week after the last self-administration session), and brains were processed for striatal tissues DA quantification with DA ELISA kit. During striatal tissues quantification, DA levels in NAc, dorsomedial striatum (DMS) and aDLS were analyzed for all rats, except for one rat where technical reasons prevented us from measuring DA levels in DMS.

#### Experiment 2

27 *TH::Cre* rats were infused with a virus coding for the inhibitory DREADD (designer receptor exclusively activated by designer drug(19)) hM4Di-mCherry or mCherry alone in the substantia nigra pars compacta (SNc). Microdialysis experiments were performed at least two weeks after the surgeries to allow the proper recovery of the animals and a stable transgene expression (20). At the end of the experiment, rats were euthanized, and brains were processed for histological validation of the probe placement and transgenes expression.

#### Experiment 3

Similarly to experiment 1, 25 *TH::Cre* rats were trained in IA 20%-EtOH 2BC and operant ethanol self-administration. After 38 sessions of operant ethanol self-administration, viral surgeries were performed. Following two weeks of recovery, hM4Di and mCherry rats were trained for 8 additional sessions, allowing stabilization of performances, before starting footshock-punished ethanol self-administration. At this step, 5 punished sessions under saline treatment were conducted to identify rats without compulsive alcohol-related behaviors (17 rats), followed by 6 unpunished sessions in absence of treatment (wash period) and, finally, 6 punished sessions under DREADDs agonist Compound 21 (C21) treatment. At the end of the experimental procedure, the sensitivity to footshocks was measured. Rats were then euthanized, and brains were processed for histological validation of transgenes expression.

#### Experiment 4

48 *TH::Cre* rats were trained in operant sucrose self-administration during 15 sessions before viral surgeries. After 14 days of recovery, hM4Di and mCherry rats were trained again during 5 sessions, allowing stabilization of performances (≤ 30% performance variation over three consecutive sessions). Then, they were tested for 3 sessions under C21 or saline treatment. Finally, recovery from treatments (wash period) was assessed during 6 sessions. Following sucrose self-administration experiment, rats performed in this following order: 2.5% sucrose two-bottle choice drinking test, open arena test, stepping test, light/dark avoidance test, elevated plus-maze and forced-swim test. Over the 48 rats, 11 rats that did not reach a minimum of 40 rewards obtained during a session were excluded of from this protocol but kept for the rest of the tests. At the end of the experimental procedure, rats were euthanized, and brains were processed for histological validation of transgenes expression.

### hM4Di-DREADDs expression

Chemogenetic manipulation of SNc DA neurons was achieved through stereotaxic bilateral infusion of a AAV5-hSyn-DIO-hM4D(Gi)-mCherry (10^12^ particles/mL, plasmid #44362, Addgene, Watertown, MA, USA) or AAV5-hSyn-DIO-mCherry (10^12^ particles/mL, plasmid #50459, Addgene) in the SNc of *TH::Cre* rats (20,21). As previously described and validated in (20), 1 μL was infused at a rate of 0.2 μL/min on each hemisphere and coordinates for SNc injections were set at: -4.3 mm (AP), ±2.4 mm (ML), -7.9 mm (DV) relative to bregma (22).

### hM4Di-DREADDs activation

DREADD agonist C21 (Hello Bio, Bristol, UK) was dissolved in 0.9% saline and kept at -20°C before testing. All the injections were given intraperitoneally at a dose of 0.5 mg/kg (1 ml/kg bodyweight). Saline solution (NaCl 0.9%, Sigma, Saint-Quentin-Fallavier, France) was prepared and kept in identical conditions.

### *In vivo* microdialysis experiment

Microdialysis experiment was carried out as previously described in (23) and is deeply detailed in Supplement 1. Briefly, homemade microdialysis probes were prepared and bilaterally implanted in the NAc and aDLS. The stereotaxic coordinates were set at: +2.2 mm (AP); ±3.2 mm (ML) and -6.5 mm (DV) for the aDLS; +2.6 mm (AP); ±1.2 mm (ML) and -8 mm (DV) for the NAc, relative to bregma (22). The dialysis fractions were collected every 45 min, over a 6-h period divided in a 1h30 period without treatment and a 4h30 period following injection of C21 or saline solution. After histological validation, DA contents of dialysis fractions were determined using high-performance liquid chromatography (HPLC).

### Behavioral procedures

#### Intermittent access to 20% alcohol two-bottle choice drinking procedure

High levels of voluntary alcohol consumption were obtained in IA 20%-EtOH 2BC drinking procedure as previously described (24,25). Briefly, single-housed rats were given 24h concurrent access to one bottle of 20% (v/v) alcohol (VWR International S.A.S., Fontenay-sous-Bois, France) in tap water and one bottle of water with 24 or 48h alcohol deprivation periods between the alcohol-drinking sessions. At the end of the procedure, rats consuming less than 2 g/kg/24h over the last three days were excluded from the study.

#### Operant ethanol self-administration

Rats were trained to orally self-administer the 20% ethanol solution in operant self-administration chambers housed within light-resistant, sound-attenuating boxes (ENV-022MD, Med Associates, St Albans, VT, USA). At the beginning of each session, two levers were extended around the liquid cup: an active, reinforced, lever, for which presses resulted in the delivery of 0.1 ml of the ethanol solution (20% v/v) in the liquid cup associated to light stimulus above the lever, and an inactive, non-reinforced, lever for which a press causes neither delivery of ethanol nor light stimulus. Lever presses on both lever and reinforcer deliveries were recorded by MED-PC IV software (Med Associates). After 2 to 3 nights in the chambers to allow acquisition of a lever-press response for ethanol under a fixed ratio 1 (FR1), operant sessions were conducted 5 days per week, with the schedule requirement increased to FR3 and the length of session shortened from 60 to 30 min over the first 2 weeks (25). Rats pressing for less than 0.3 g/kg/30min at the end of the training were excluded from the study.

#### Footshock-punished ethanol self-administration

Identification of rats expressing compulsive alcohol-related behavior, defined by the persistence of alcohol use despite knowledge of negative associated consequences (3), was achieved by coupling ethanol self-administration with intermittent footshocks. During punished sessions, self-administration parameters were identical to baseline self-administration (i.e., FR3, 30 min) but footshocks (0.25 mA, 0.5 s) were delivered each 8 presses on the reinforced lever, following the paradigm developed by (26). By doing so, footshock punishment coincide with the preparatory response (lever presses) but not consummatory behaviors (alcohol delivery), and favor the punishment of seeking rather than of taking (18,27). Rats that continued to press the lever, despite footshocks (≥ 70% of the baseline performances), were considered as footshock-resistant whereas rats that stop to press the lever in face of the negative consequence were considered as footshock-sensitive. This cut-off was determined based on the variances of the basal operant performances we standardly obtained and the bimodal distribution of our rats’ population (see also Supplementary Fig. S1A in Supplemental 1).

#### Sensitivity to footshocks

Rats were place in a StartFear chamber (Panlab, Barcelona, Spain) and footshocks were delivered at 0.2, 0.25 and 0.3 mA. At each intensity, triplicates were made with 20 s spaced, and 1 min was left before increasing the intensity. Footshocks threshold was defined as a freeze or a jump off the grid. Footshock responses were recorded with a video-tracking system (EthoVision XT 15 software, Noldus Information Technology) and scored by two observers blind to the experimental conditions.

#### Operant sucrose self-administration

Rats were trained to orally self-administer a 2.5% sucrose (Sigma, Saint-Quentin-Fallavier, France) solution as described for the operant ethanol self-administration, except that each lever press results in the delivery of 0.2 mL of sucrose (13). After 2 to 3 nights in the chambers to allow acquisition of a lever-press response for sucrose under a fixed ratio 1 (FR1), 60 min operant sessions were daily conducted with a maximum of sucrose deliveries set at 100 before ending the session.

#### 2.5% sucrose two-bottle choice drinking test, open arena test, stepping test, light/dark avoidance test, elevated plus-maze and forced-swim test

They were performed according to (13). A fully detailed description of the methodological procedures is provided for each behavioral test in Supplement 1.

### Striatal tissues DA quantification

aDLS, DMS and NAc were dissected from frozen brains and homogenized in: HCl 0.01N, EDTA 1 mM and Na2S2O5 4 mM to prevent catecholamine degradation. DA levels were next measured using DA ELISA kit (Immusmol SAS, Talence, France) according to the manufacturer’s instruction and the literature (28,29). At the end of the reaction, a sulfuric acid 0.25 M was added, and the optical density was detected by spectrophotometer (PHERAstar, BMG Labtech, Champigny sur Marne, France) at the wavelength of 450 nm. A reference wavelength was additionally used at 620 nm to evaluate non-specific absorbance. Finally, DA levels were determined by comparing absorbance of samples with external standards and were expressed as the amount of DA, in ng per mg of protein present in the homogenate determined with BCA Protein Assay Kit (Thermo Scientific, Illkirch, France).

### Histological analysis

#### Brain tissue preparation and processing

A detailed description of brain tissue preparation and processing is provided in Supplement 1.

#### Quantification of transgenes expression

To assess transgenes expression in DA mesencephalic regions, immunostaining for tyrosine hydroxylase (TH) were carried out as previously described (20) and as detailed in Supplement 1. DREADD expression was quantified for each hemisphere by comparing the number of TH-labeled-mCherry-positive neurons with the number of TH-labeled neurons within three areas: the distal SNc, the medial SNc and the ventral tegmental area. Rats with less than 30% of transgene expression in the distal SNc in each hemisphere were excluded from the study (20).

### Statistical analyses

Statistical analyses are described in figure legends, summarized in Table S1 and fully detailed in Supplement 1.

## Results

### Compulsive alcohol use is specifically associated with a decrease in DA levels in the aDLS

We first tested whether compulsive alcohol use is associated with a hypodopaminergic state of the nigrostriatal pathway, by assessing striatal DA levels in rats expressing, or not, compulsive alcohol seeking behavior. In the first experiment, animals were first trained to voluntary consume IA 20%-EtOH 2BC, before being challenging instrumentally to respond for alcohol in operant chambers (24,25). When operant performances reached stable levels, lever presses for alcohol were coupled to mild footshocks to identify rats expressing compulsive alcohol use, operationalized as resistance to punishment of alcohol self-administration (26,30) (Fig. 1A). During these punished sessions, a bimodal distribution progressively appeared, which clearly indicates the existence of two distinct sub-populations: one with a strong decrease in lever presses, the footshock-sensitive (FS) rats (58%), while the other, the footshock-resistant (FR) rats (42%), maintained their responses despite footshocks (Fig. 1, C and D, Supplementary Fig. S1A in Supplement 1; see also (30,31)). No differences were observed on the second, inactive lever, indicating that the tendency to resist to the punishment was specifically related to the search for alcohol (Supplementary Fig. S1B in Supplement 1). Importantly, and as already observed for alcohol (18) and cocaine (32), the history of intake cannot account for these two distinct phenotypes. Indeed, we found a similar escalation of alcohol consumption during IA 20%-EtOH 2BC (Fig. 1B and Supplementary Fig. S1C in Supplement 1), as well as similar lever presses levels during unpunished self-administration sessions between FR and FS rats (Fig. 1, C and D). While DA levels measured by ELISA assays on striatal extracts did not differ between FS and FR rats in the NAc or DMS and appeared similar to those obtained from a non-exposed alcohol condition (water control rats), FR rats showed a significant decrease in DA levels in the aDLS (Fig. 1E). In addition, a robust negative correlation was found between the degree of resistance to footshocks and the amount of aDLS DA (Fig. 1F). Thus, in line with our working hypothesis, compulsive alcohol seeking is associated with a decrease in aDLS DA level, suggesting an important link between a tonic hypodopaminergic state of the nigrostriatal pathway and the emergence of this maladaptive behavior.

**Fig. 1:**
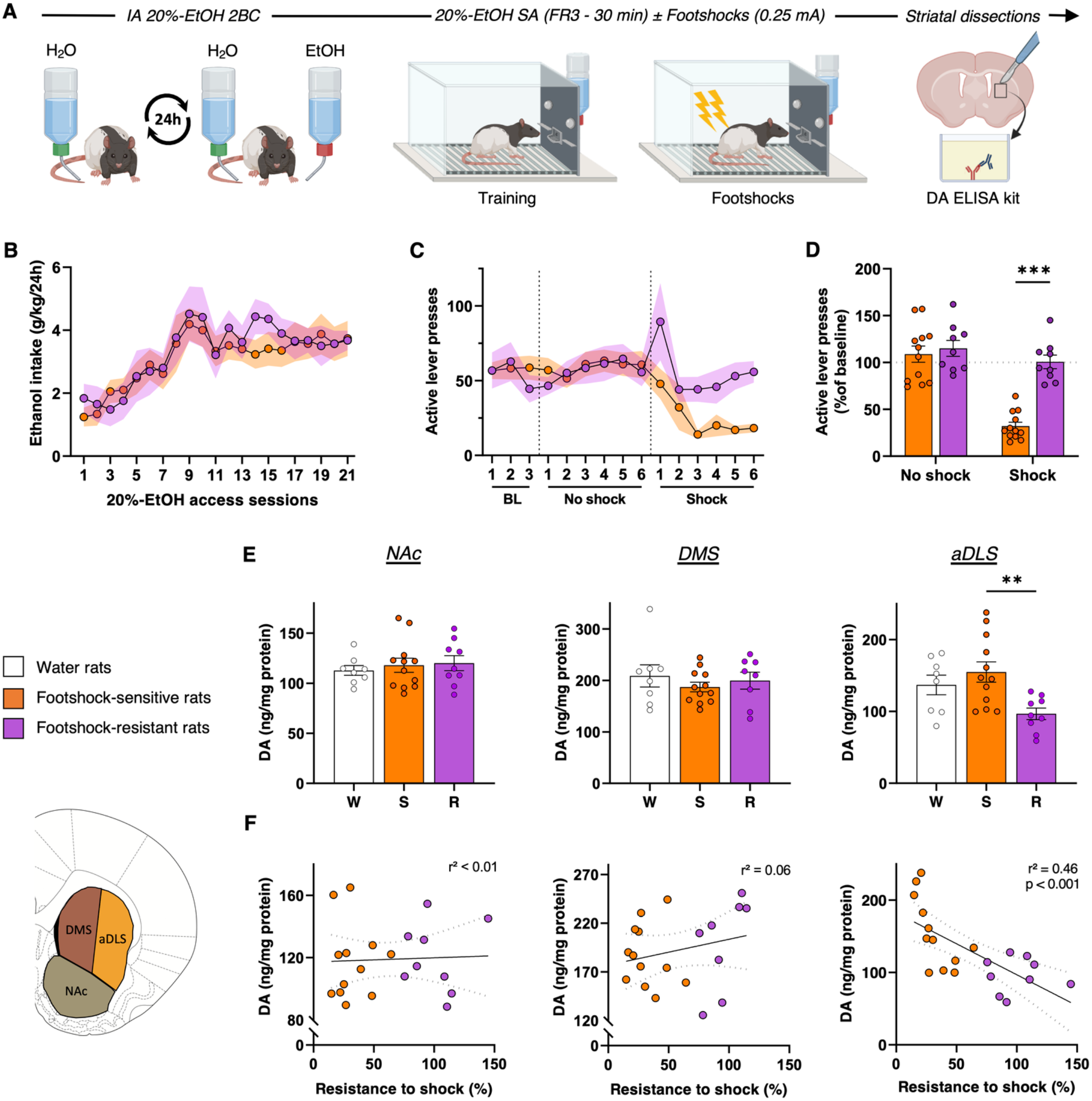
Compulsive alcohol use is specifically associated with a decrease in DA levels in the aDLS. **(A)** Experiment timeline. **(B)** Ethanol intake during intermittent-access 20%-ethanol two-bottle-choice (*IA 20%-EtOH 2BC*). RM two-way ANOVA showed a significant effect of session [F_(6, 109)_=7.82, P<0.001, partial η^2^=0.29] but neither effect of group, nor session x group interaction [F_s_<0.47, P>0.5, partial η^2^<0.02]. **(C)** Number of active lever presses in 30-min self-administration sessions (SA) of 20% EtOH (FR3), during baseline, “no-shock” and “shock” sessions (see supplementary material for details) in *footshock-sensitive* (FS, orange, *n*=12) and *footshock-resistant* (FR, purple, *n*=9) rats. RM two-way ANOVA showed a significant group x session interaction [F_(14, 266)_=2.86, P<0.001, partial η^2^=0.13]. **(D)** Mean active lever presses during the three last “no-shock” sessions and the three last “shock” sessions normalized to baseline. RM two-way ANOVA showed a significant shock condition x group interaction [F_(1, 19)_=20.89, P<0.001, partial η^2^=0.52] **(E)** NAc, DMS and aDLS DA levels for *FR, FS* and *Water* rats (white, *n*=8). One-way ANOVA showed a main effect of group in the aDLS [F_(2,26)_=5.56, P<0.01, partial η^2^=0.3], but not in the NAc and DMS [F_s_<0.55, P>0.5, partial η^2^<0.04]. **(F)** Linear regression between resistance to footshocks and DA level in NAc, DMS or aDLS. A significant correlation was found in the aDLS [F_(1,19)_=16.18, P<0.001], but not in the NAc or DMS [F_s_<1.14, P>0.3]. Data are expressed in mean ± SEM. Bonferroni correction post-hoc analysis: **, P<0.01; ***, P<0.001. BL: baseline, DMS: dorsomedial striatum, aDLS: anterior dorsolateral striatum, FR: fixed ratio, NAc: nucleus accumbens.

### Chemogenetically-induced nigrostriatal hypodopaminergia induces compulsive alcohol use

Consequently, we next investigated whether this DA hypofunction is causally implicated in the emergence of compulsive alcohol use. To that end, an experimental and reversible nigrostriatal hypodopaminergia was induced using the inhibitory DREADD hM4Di (19). *TH::Cre* rats were transduced in the SNc with AAVs that Cre-dependently express hM4Di-mCherry, or mCherry alone (control condition), allowing its specific expression in SNc DA neurons (Fig. 2, A-C). hM4Di was activated by the synthetic ligand C21 (33), at a dose that we previously reported as potent and selective to inhibit SNc neurons in *TH::Cre* rats (20). We first confirmed with *in vivo* microdialysis that this approach efficiently decreases aDLS DA levels (Fig. 2D and Supplementary Fig. S2 in Supplement 1). In hM4Di rats, a 30%-decrease in aDLS DA level was observed between 90 and 135 minutes after C21 injection (Fig. 2E), which is consistent with the temporal inhibition of SNc neurons previously reported (20). Importantly, this strategy did not influence NAc DA levels (Fig. 2E), confirming that chemogenetic inhibition of SNc DA neurons induces a reversible and selective hypodopaminergic state of the nigrostriatal pathway.

**Fig. 2:**
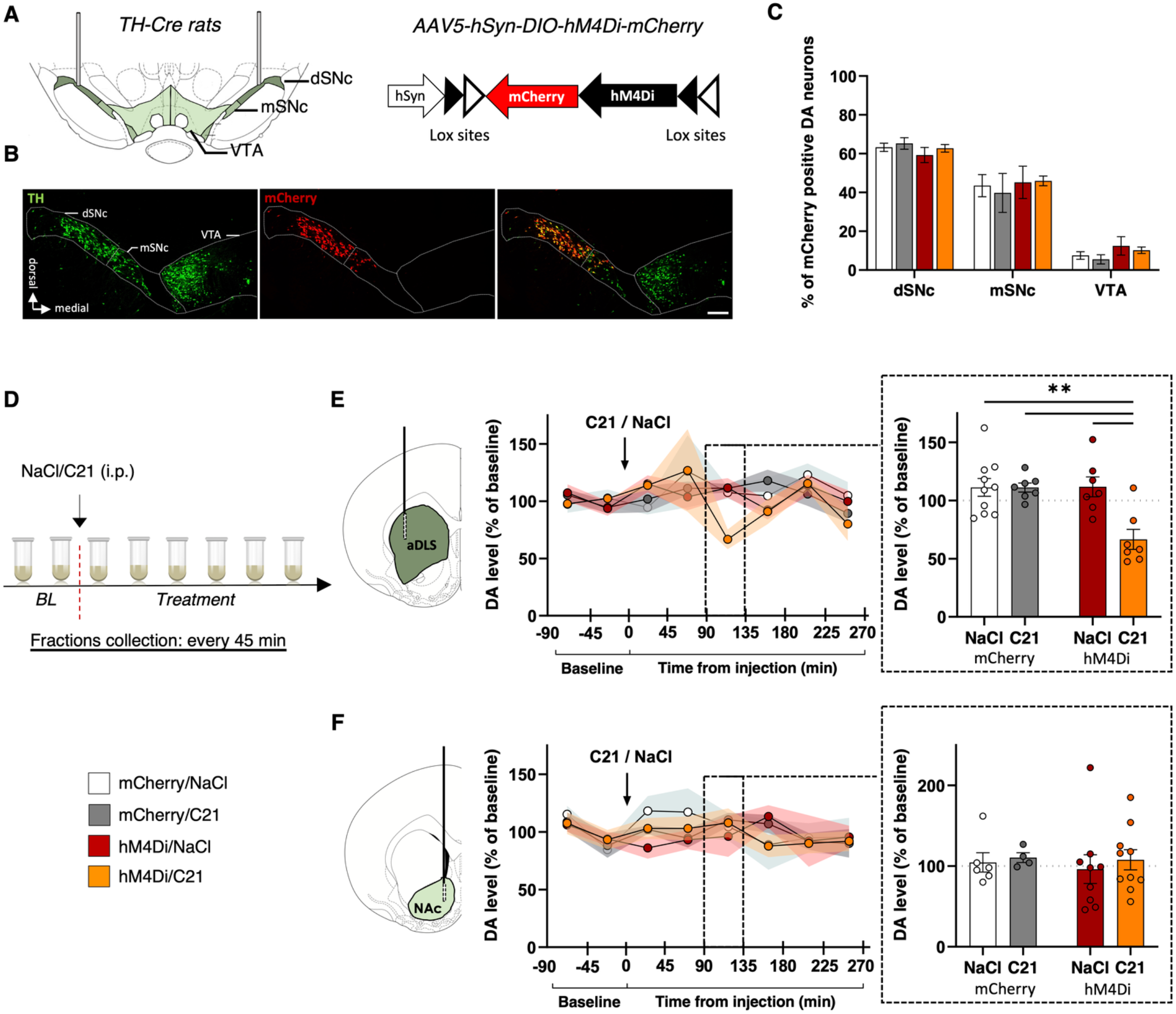
Chemogenetic inhibition of the SNc DA neurons induces selective nigrostriatal hypodopaminergia. **(A-C)** hM4Di-mCherry and mCherry expression in SNc DA neurons. **(A)** *TH::Cre* rats received bilateral injection of Cre-dependent hM4Di-mCherry or Cre-dependent mCherry virus in the SNc. **(B)** Representative images of TH immunostaining and hM4Di-mCherry expression. Scale bar: 250 μm. **(C)** Quantification of transgene expression in distal (dSNc), medial SNc (mSNc) and VTA. Three-way ANOVA showed a significant effect of the area [F_(2,129)_=158, P<0.001, η^2^=0.71], but no effect of transgene, treatment or any interaction between these factors [F_s_<1.04, P>0.36, partial η^2^<0.02]. **(D-G)** Extracellular DA concentrations in aDLS and NAc throughout eight 45 min-fractions collected by microdialysis. Data are normalized to baseline. **(D)** Design of the microdialysis experiment. **(E)** Time course of extracellular DA concentrations in the aDLS of *hM4Di* rats treated with C21 (orange, *n*=7) or NaCl (red, *n*=7) and *mCherry* rats treated with C21 (grey, *n*=7) or NaCl (white, *n*=10). **(F)** Time course of extracellular DA concentrations in the NAc of *hM4Di* rats treated with C21 (orange, *n*=10) or NaCl (red, *n*=9) and *mCherry* rats treated with C21 (grey, *n*=4) or NaCl (white, *n*=6). In the fraction collected between 90 and 135 minutes after injection (dotted squares), two-way ANOVA found a treatment x transgene interaction in the aDLS [F_(1,27)_=8.63, P<0.01, partial η^2^=0.24], but not in the NAc [F_(1,25)_=0.03, P=0.87, partial η^2^=0.001]. Data are expressed in mean ± SEM. Bonferroni correction post-hoc analysis: **, P<0.01. C21: compound 21, SNc: substantia nigra pars compacta, TH: tyrosine hydroxylase, VTA: ventral tegmental area.

Then, we tested whether induction of this experimental nigrostriatal hypodopaminergia state is sufficient to induce compulsive alcohol seeking in FS rats. In the third experiment, *TH::Cre* rats were therefore trained in IA 20%-EtOH 2BC and operant ethanol-self-administration procedures as in the first experiment (Fig. 3A), and then transduced with DREADDs ensuring a same history of alcohol exposure between hM4Di and mCherry control rats (Fig. 3, B and C). After selection of FS-hM4Di and FS-mCherry rats under punishment sessions preceded by saline injections, we assessed their tendency to persist in seeking alcohol despite punishment over the course of a second series of sessions, this time following administration of C21 (Fig. 3D). In these sessions, we found that chemogenetic inhibition of SNc DA neurons progressively increased the resistance of FS rats to punishment (Fig. 3, D and E), while it did not affect their intrinsic footshocks sensitivity (Supplementary Fig. S3A in Supplement 1). Interestingly, this increase was observed for each hM4Di FS rat (Fig. 3F) and correlated only with the level of hM4Di expression within the distal SNc (Fig. 3, G and H), the main DA input of the DLS (34). In marked contrast, effect of the chemogenetic manipulation was found neither on the inactive lever during punishment sessions (Supplementary Fig. S3B in Supplement 1), nor on operant alcohol self-administration under baseline conditions (Supplementary Fig. S3C in Supplement 1), confirming that this effect was related to punishment and not to a non-selective or general change in operant behavior. Taken together, these results demonstrated that an hypodopaminergic state of the nigrostriatal pathway is sufficient to induce compulsive alcohol seeking behavior in animals that were otherwise resilient.

**Fig. 3:**
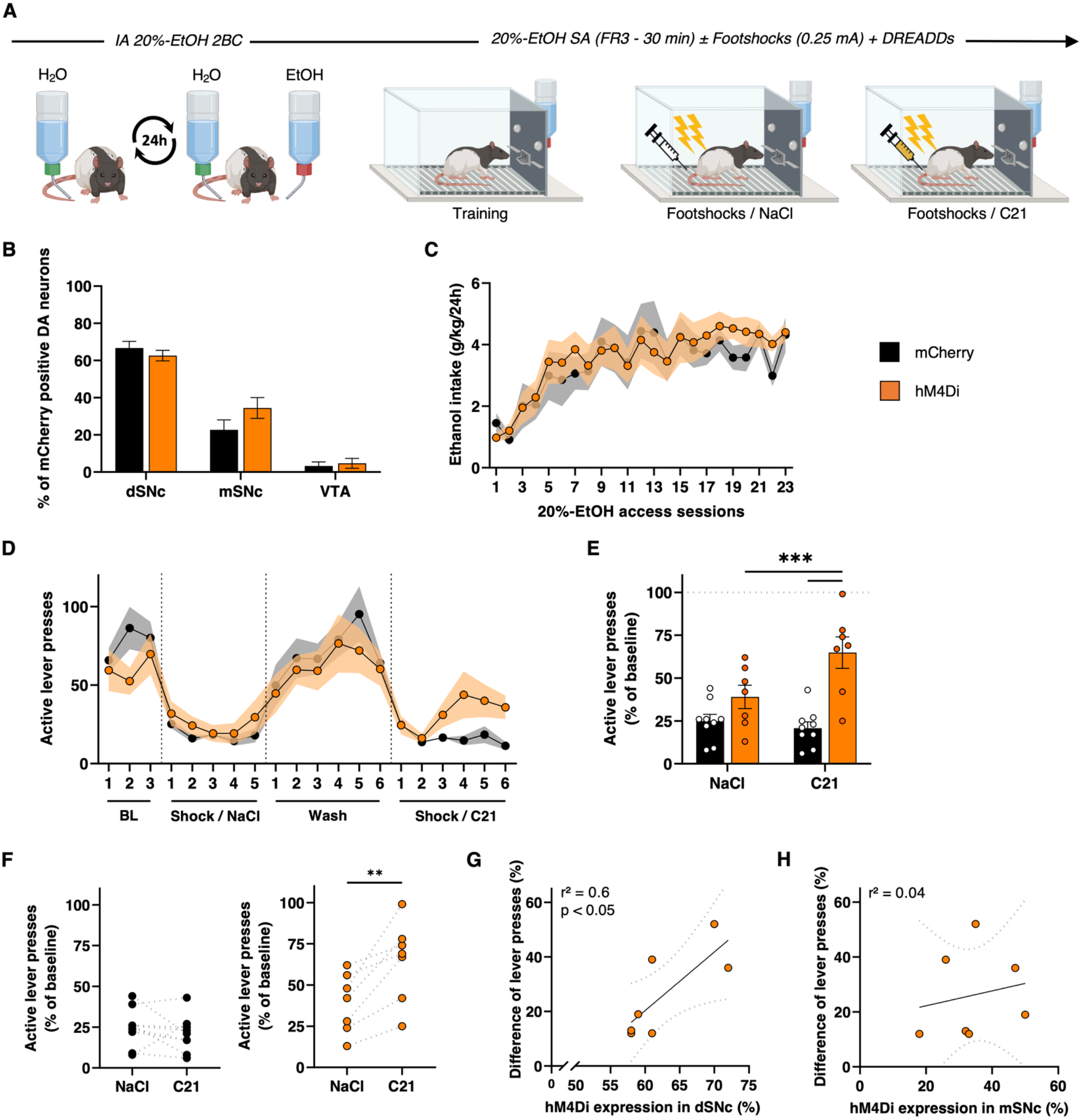
Chemogenetically-induced nigrostriatal hypodopaminergia induces compulsive alcohol use. **(A)** Experiment timeline. **(B)** Quantification of hM4Di-mCherry and mCherry expression in distal (dSNc), medial SNc (mSNc) and VTA. Two-way ANOVA showed significant effect of the area [F_(2,90)_=116.4, P<0.001, η^2^=0.72], but no effect of transgene, treatment or any interaction between these factors [F_s_<1.99, P>0.14, partial η^2^<0.04]. **(C)** Ethanol intake during intermittent-access 20%-ethanol two-bottle-choice (*IA2BC 20%-EtOH*) in *hM4Di* rats (orange, *n*=7) and *mCherry* rats (black, *n*=10). RM two-way ANOVA showed a significant effect of session [F_(4, 59)_=10.56, P<0.001, partial η^2^=0.43), but neither effect of transgene, nor session x transgene interaction [F_s_<0.57, P>0.5, partial η^2^<0.04]. **(D)** Number of active lever presses in 30-min self-administration sessions of 20%-EtOH (FR3), during baseline, “Shock/NaCl”, “Wash” and “Shock/C21” sessions. RM two-way ANOVA showed a significant session x transgene interaction [F_(19,266)_=2.18, P<0.01, partial η^2^=0.13]. **(E - F)** Mean active lever presses during the three last “Shock/NaCl” and “Shock/C21” sessions normalized to baseline **(E)** and individual trajectories during these two periods **(F)**. RM two-way ANOVA reported a significant session x transgene interaction [F_(1,14)_=18.41, P<0.001, partial η^2^=0.57], while paired t-test reported a significant effect of C21 in *hM4Di* [t=4.3, P<0.01] but not in *mCherry* rats [t=1.02, P=0.34]. **(G - H)** Correlation between the difference of active lever presses during “Shock/C21” and “Shock/NaCl” sessions and the percent of hM4Di expression within dSNc **(G)** or mSNc **(H)**. A significant correlation was found for dSNc [F_(1,5)_=7.64, P<0.05], but not for mSNc [F_(1,5)_=0.18, P=0.69]. Data are expressed in mean ± SEM. Bonferroni correction post-hoc analysis: **, P<0.01; ***, P<0.001.

### Chemogenetically-induced nigrostriatal hypodopaminergia leads to a prolonged negative motivational, but not affective, state

Because we previously showed that a partial DA denervation of the nigrostriatal pathway leads to motivational impairments as well as depression- and anxiety-related behaviors (13), three core features of the negative psychological state experienced during withdrawal (35), we finally test whether chemogenetically-induced nigrostriatal hypodopaminergia recapitulates such phenotype (Fig. 4, A and B). Motivated behavior was evaluated in an operant sucrose self-administration task as previously performed (13,14). In comparison to the control groups, hM4Di-C21 treated rats exhibited a prolonged decrease in their performance to obtain sucrose, that persisted beyond the last session under C21 (Fig. 4, C and D). This result was not due to a motor deficit associated with nigrostriatal hypodopaminergia, as revealed by the absence of motor alterations in an open arena and fine use of the forepaws in a stepping test (Fig. 4, E and F). It cannot be attributed either to an inability to discriminate between the active and inactive lever that was preserved throughout the task (Supplementary Fig. S4A in Supplement 1), or to a decrease in sensitivity to the reinforcing properties of sucrose, as preference (Fig. 4G) and consumption (Supplementary Fig. S4B, in Supplement 1) for the sucrose solution was unchanged in a two-bottle choice paradigm. Taken together, these data confirm that the chemogenetically-induced decrease in operant sucrose behavior reflects a motivational deficit (13). In sharp contrast, no increase in anxiety- or depression-related behaviors were observed in light/dark avoidance and elevated-plus maze tests (Fig 4. H and Supplementary Fig. S4C in Supplement 1) and in the forced swim test (Fig. 4I) respectively. Finally, in addition to its implication in compulsive alcohol use, hypodopaminergic state of the nigrostriatal pathway leads to a prolonged negative motivational, but not affective, state.

**Fig. 4:**
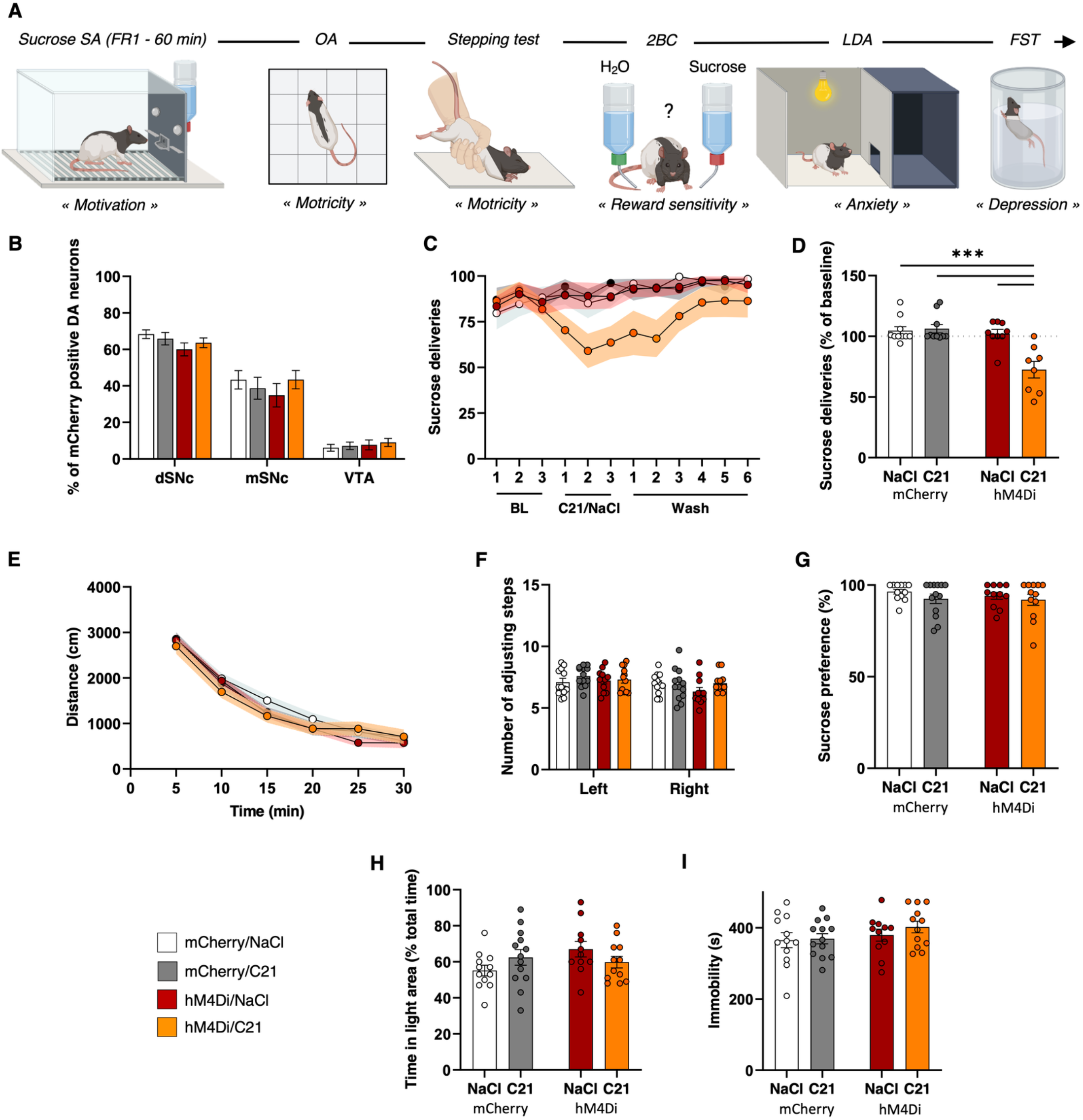
Chemogenetically-induced nigrostriatal hypodopaminergia leads to a prolonged negative motivational, but not affective, state. **(A)** Behavioral screening timeline. **(B)** Quantification of hM4Di-mCherry and mCherry expression in distal (dSNc), medial SNc (mSNc) and VTA. Three-way ANOVA showed significant effect of the area [F_(2, 140)_=346.7, P<0.01, η^2^=0.83], but no effect of transgene, treatment or any interaction between these factors [F_s_<1.31, P>0.27, partial η^2^<0.02]. **(C)** Number of 2.5%-sucrose deliveries obtained in 60-min self-administration (SA) sessions under continuous reinforcement (FR1), during baseline, “C21/NaCl” and “Wash” sessions, in *hM4Di* rats treated with C21 (orange, *n*=8) or NaCl (red, *n*=9) and *mCherry* rats treated with C21 (grey, *n*=10) or NaCl (white, *n*=10). RM three-way ANOVA showed a significant session x transgene x treatment interaction [F_(11, 363)_=2.67, P<0.01, partial η^2^=0.07]. **(D)** Mean sucrose deliveries obtained during “C21/NaCl” sessions normalized to baseline. Two-way ANOVA showed a significant transgene x treatment interaction [F_(1, 33)_=13.38, P<0.001, partial η^2^=0.29]. **(E - I)** Distance traveled over the course of a 30-min session in an open area (OA) **(E)**, number of adjusting left and right forepaws in a stepping test **(F)**, sucrose preference measured over a 60-min 2.5%-sucrose two-bottle-choice (2BC) drinking session **(G)**, percentage of time spent in the light area in a light/dark avoidance test (LDA) **(H)** and time spent immobile in a forced swim test (FST) **(I)** in *hM4Di* rats treated with C21 (orange, *n*=12) or NaCl (red, *n*=11) and *mCherry* rats treated with C21 (grey, *n*=13) or NaCl (white, *n*=12). Two- or three-way ANOVA found no interaction implicating the transgene and the treatment [F_s_<1.3, P>0.26, partial η^2^<0.03] in these different tests, except a marginal side x transgene x treatment interaction in stepping test [F_(1, 44)_=3.55, P=0.07, partial η^2^=0.07] and a transgene x treatment interaction in LDA [F_(1, 44)_=3.68, P=0.06, partial η^2^=0.08], mainly driven by small size effects related to the transgene condition and not to a C21 effect on *hM4Di* rats. Data are expressed in mean ± SEM. Bonferroni correction post-hoc analysis: ***, P<0.001.

## Discussion

At a time of great controversies about the exact role of DA in addiction (12), we shed to light here hitherto overlooked implication of the DA nigrostriatal pathway in AUD. Our findings indicate that chronic exposure to alcohol leads to the development of a tonic hypodopaminergic state in the nigrostriatal pathway in vulnerable individuals that directly contributes to the development of compulsive alcohol use and probably participates to the maintenance of the negative motivational state experienced during abstinence.

Mesolimbic tonic hypodopaminergic states have been proposed to be part of the allostasis phenomenon observed during abstinence that underlines the emergence of a negative motivational and affective state and participates to the development of excessive alcohol seeking and intake (9,36). In regard to the present data, it suggests that, in vulnerable chronic users, these DA hypofunctions might eventually propagate or migrate to the nigrostriatal pathway through different possible interconnexions (37,38). This might be the tonic counterpart of striatal DA phasic signals transition, that have been observed to emerge progressively in the aDLS over the course of drug taking, while they concomitantly fade in the ventral portion (39). Interestingly, DA phasic signals are directly modulated by DA tonic state (40) and the relation between these two complementary modes of signaling has been suggested to be important to consider for alcohol and other substance use disorder (41). Emergence of the hypodopaminergic state of the mesolimbic pathway in the early phase of the disorder might therefore lead to an aberrant enhanced DA phasic signaling in the NAc, that would contribute to aberrant learning (42) and sensitization of the mesolimbic DA pathway to drugs or associated cue (43,44). Within this framework, optogenetic studies conducted within the mesolimbic pathway have shown that increasing DA phasic firing promotes excessive alcohol seeking, whereas increasing DA tonic firing decrease consumption and annihilates the effect of DA phasic signal when induced simultaneously (45,46). In the ultimate stages of AUD, the hypodopaminergic state of the nigrostriatal pathway revealed in our study, would strengthen similar abnormal phasic DA signals in the aDLS engaged by drug-paired cues, facilitating the formation of maladaptive incentive habits and the ensuing rigidity in drug seeking (18,38,47) exacerbated by negative urgency during abstinence (48). This therefore suggests that DA-based therapy normalizing DA tonic firing may eliminate aberrant DA phasic signals and suppress excessive as well as compulsive alcohol seeking and drinking. In this respect, normalization of DA tone in the mesolimbic pathway, using GDNF or OSU6162, has already been shown effective at suppress excessive alcohol consumption in rodents and diminish alcohol craving in humans (9,49,50). Further studies, build on the present findings, would be of high interest to determine whether normalization of tonic DA state in the nigrostriatal pathway with new DA-based therapy is efficient to limit compulsive alcohol use.

Beside compulsivity, our study also intriguingly reports a dissimilar effect of the nigrostriatal hypodopaminergia on the motivation to obtain alcohol and sucrose. Indeed, whereas nigrostriatal hypodopaminergia did not change motivation to self-administer alcohol, and even increased resilience to obtain alcohol in the presence of footshocks, it markedly decreases motivation to self-administer sucrose. It suggests that hypodopaminergic state of the nigrostriatal pathway may have different behavioral outcomes depending on the history associated to- and the natural properties of the reward. If this observation requires further investigations to directly compare motivation for sucrose and alcohol (51), it represents an interesting feature to note, as a shift in interest in the drug at the expense of other natural rewards or activities is an important dimension of addiction (2,52,53).

In conclusion, this study point toward the causal implication of a tonic hypodopaminergic state of the nigrostriatal pathway in compulsive alcohol use, a cardinal feature of AUD (3). These data also emphasize the importance of the nigrostriatal DA pathway in motivated behavior and suggest that this hypodopaminergic state contributes to the negative motivational state observed during withdrawal. The role of this pathway in motivation-related processes has long been neglected (13), while it could represent one of the key elements to understanding the complex physiopathology of AUD. Future works will be necessary to decipher the neurobiological and molecular mechanisms underlying this tonic hypodopaminergic state, as well as how this state leads to the emergence of compulsive alcohol use. Although these questions remain, it is likely to open new avenues of research in AUD and therapeutic strategies based on the restoration of a normal DA tone to annihilate this maladaptive behavior.

## Supplementary information

Supplementary information is available at MP’s website

## Funding

This work was supported by the Institut National de la Santé et de la Recherche Médicale (Inserm), the Agence Nationale de la Recherche (ANR-16-CE16-0002, to SC) and Grenoble Alpes University.

## Acknowledgments

We thank Pr David Belin for critical reading of the manuscript; the PIC GIN Platform for technical assistance in fluorescence microscopy and analysis, as well as the GIN behavioral facility that is supported by the Grenoble Center of Excellence in Neurodegeneration (GREEN).

## Conflict of Interest

Authors report no conflict of interest.

## SUPPLEMENT 1

## SUPPLEMENTAL METHODS

### Animals

All animals used in this study were bred at the Plateforme Haute Technologie Animal or at the Grenoble Institut Neurosciences, La Tronche, France). All experimental protocols complied with the European Union 2010 Animal Welfare Act and the new French directive 2010/63, and were approved by the French national ethics committee no. 004.

### hM4Di-DREADDs expression

Chemogenetic manipulation of SNc DA neurons was achieved through stereotaxic bilateral infusion of a AAV5-hSyn-DIO-hM4D(Gi)-mCherry (10^12^ particles/mL, plasmid #44362, Addgene, Watertown, MA, USA) or AAV5-hSyn-DIO-mCherry (10^12^ particles/mL, plasmid #50459, Addgene) in the SNc of *TH-Cre* rats (1,2). Animals were anesthetized with a mixed intraperitoneal injection of ketamine (Chlorkétam, 60 mg/kg, Mérial SAS, Lyon, France) and xylazine (Rompun, 10 mg/kg, Bayer Santé, Puteaux, France). Then local anesthesia was provided by a subcutaneous injection of lidocaïne (Lurocaïne, 8 mg/kg, Vetoquinol S.A., Lure, France) on the skull surface and animal were secured in a Kopf stereotaxic frame (Phymep, Paris, France) under a microbiological safety post (PSM). Coordinates for SNc injections were determined according to (3), adjusted to the body weight, and set at, relative to bregma: -4.3 mm (AP), ±2.4 mm (ML), -7.9 mm (DV). On each hemisphere, 1 μL was infused at a rate of 0.2 μL/min using microinjection cannula (33-gauge, Plastic One, USA) connected to a 10 μL Hamilton syringe and a microinjection pump (Stoelting Co., Wood Dale, IL, USA). After each injection, the cannula was left in position for 5 min to allow the injected solution to be absorbed into the parenchyma and minimize the spread of the virus along the cannula tract. The skin was sutured, disinfected, and the animal placed in a heated wake-up cage, before being replaced in its home-cage after complete awakening and monitored for several days.

With these parameters, we always observed mCherry staining in TH-positive neurons. This selective expression of the transgene was confirmed in a pilot experiment in which no mCherry staining was detected after the total destruction of SNc DA neurons with the neurotoxin 6-hydroxydopamine (data not shown).

### *In vivo* microdialysis experiment

Rats were anesthetized by inhalation of isoflurane (Isoflurin, Axience SAS, Pantin, France) and secured in a Kopf stereotaxic frame (Phymep, Paris, France). The dorsal skull was exposed, and holes were drilled to facilitate the bilateral implantation of microdialysis probes. Homemade microdialysis probes were prepared and and the length of the dialysis membrane was adapted to the brain regions studied (1.8 mm for the anterior dorsolateral striatum (aDLS) and 1 mm for the nucleus accumbens (NAc)). During each experiment, one probe was lowered in the aDLS on one side and one probe was lowered in the NAc on the other side. The stereotaxic coordinates were determined according to (3), adjusted to the body weight, and set at, relative to bregma: +2.2 mm (AP); ±3.2 mm (ML) and -6.5 mm (DV) for the aDLS; +2.6 mm (AP); ±1.2 mm (ML) and -8 mm (DV) for the NAc. The aDLS was exclusively targeted here (see Supplementary Fig. S2), as well as in the striatal tissues DA quantification experiment, because compulsive alcohol seeking in rats has been recently associated with this territory specifically (4). After implantation, probes were equilibrated for 1 h with artificial cerebro-spinal fluid (NaCl 149 mM, KCl 2.8 mM, MgCl_2_ 1.2 mM, CaCl_2_ 1.2 mM, and glucose 5.4 mM, pH 7.3) at a flow rate of 1 μl/min. Then, dialysis fractions were collected with a refrigerated autosampler (820 Microsampler, Univentor, Zejton, Malta) every 45 min, over a 6-h period divided in a 1h30 period without treatment and a 4h30 period following the injections. At the end of the experiment, the skin was stitched, disinfected, and the rat placed in a heated wake-up cage, before being replaced in its home-cage after complete awakening. At least one week after the experiment, microdialysis was performed a second time by reversing the dialyzed structures on each hemisphere. After histological validation, DA contents of dialysis fractions were determined using high-performance liquid chromatography (1200 series, AGILENT Technologies, USA), with electrochemical detection (Dionex UltiMate 3000, Thermo Scientific, Illkirch, France) and an Aquasil C18 reverse-phase microcolumn (RP-18, 100 × 2.1 mm, 3 µ particle size, Thermo Scientific, Illkirch, France) maintained at 24°C. The mobile phase (NaH_2_PO_4_ 50 mM, EDTA 0.1 mM, sodium octyl sulfate 1.7 mM, KCl 4.5mM and 5% acetonitrile (vol/vol), adjusted to pH 3.1) was run at a flow rate of 0.4 ml/min. The working electrode potential was +550 mV and the detector sensitivity 2 nM for DA. The running time for each determination was 12 minutes. Chromatograms were collected and treated with integration software (Chromeleon 7 CDS Software, Thermo Scientific). DA concentrations were determined by comparing DA peaks with external standards and were expressed in nanomolar.

### Behavioral procedures

#### Intermittent access to 20% alcohol two-bottle choice drinking procedure

Concurrent access was given on Sunday, Tuesday, and Thursday. The placement (left or right) of each solution was alternated between each session to control for side preference, and a bottle of water was placed in a cage without rats to evaluate the spillage that was always ≤ 1 mL (< 3.5% of the total fluid intake). Level of alcohol and water consumption (g/kg/time) were measured after the first 30 min, allowing investigation of binge-related drinking behavior (*18*), and at the end of the 24h-session.

#### 2.5% Sucrose two-bottle choice drinking procedure

Rats were given 1 h concurrent access to one bottle of water and one bottle of 2.5% (v/v) sucrose. At the end of the session, the volumes of sucrose solution and water consumed were measured to determine sucrose, water and total fluid intake (mL/kg), and preference for sucrose over water (sucrose intake/total intake, expressed as a percentage).

#### Open arena test

Rats were placed in an opaque open arena apparatus (50 × 50 × 40 cm). Over a 30-min period, horizontal distance traveled (cm) was recorded and analyzed with a video-tracking system.

#### Stepping test

Rats, maintained by the posterior third of their body, were moved over a length of 90 cm by a rectilinear and regular movement from left to right and inversely along a smooth-surfaced table (5). The test was carried out in triplicates and the number of adjustments of right and left paws during displacements was counted by two observers blind to the experimental conditions.

#### Light / Dark avoidance test

The apparatus was composed of a light (50 × 40 × 40 cm) and a dark chamber (24 × 40 × 40 cm) separated by an opaque wall with a small aperture (14,5 × 7 cm), to allow rats to move freely between the chambers. The light chamber was opened at the top and lit with a white incandescent light (> 400 lux) located 70 cm above the floor of the chamber. By contrast, the dark chamber was closed at the top and was unlit (< 5 lux). Rats were placed in the center of the light chamber and the total time spent in the light and the dark chambers, over a 5-min period, were recorded with a video-tracking system and analyzed by an observer blind to the experimental conditions.

#### Elevated plus-maze test

The elevated plus-maze was composed of two opposing open arms (50 × 10 cm) and two opposing arms enclosed by high opaque walls (50 × 10 × 40 cm) suspended 55 cm above the floor. Rats were placed in the center of the elevated plus-maze and times spent in the open and closed arms, over a 5-min period, were recorded and analyzed with a video-tracking system.

#### Forced swim test

Rats were placed in a transparent cylinder of 40 cm high and 20 cm in diameter, filled with water (24 ± 1 °C) to a depth of 30 cm, for 10 minutes (5). Animal activity was recorded and analyzed with a video-tracking system.

### Histological analysis

#### Brain tissue preparation and processing

For striatal tissues DA quantification, rats were deeply anesthetized by isoflurane saturation and sacrificed by decapitation. Brains were immediately frozen in liquid nitrogen and stored at -80 °C. For the other experiments, rats were deeply anesthetized by isoflurane saturation or by injection of exagon (Axience SAS, Pantin, France) and transcardially perfused with 0.9% saline followed by 4% paraformaldehyde (PFA) in phosphate-buffered saline (PBS). They were then cryoprotected in 20% sucrose/PB for 24 h and frozen in isopentane cooled to -50°C on dry ice. Coronal sections of striatum and mesencephalon were cut using a cryostat (HM525, Microm, Francheville, France). For striatal tissues DA quantification, aDLS, DMS NAc were dissected from multiple thick sections of striatum (100 µm, 2.2 to 0.7 mm anterior to bregma) and stored at -80 °C before being homogenized. To verify the placement of the microinjection cannula and microdialysis probes, brains were cut at a thickness of 30 µm. Sections were then stained using Cresyl violet staining and visualized under a light microscope (Nikon Eclipse 80i, TRIBVN, Châtillon, France) coupled to the ICS FrameWork computerized image analysis system (Calopix 2.9.2 software, TRIBVN, Châtillon, France). Finally, to quantified transgenes expression, floating coronal sections (30 µm) of three levels of the mesencephalon were selected according to (1).

#### Quantification of transgenes expression

Free-floating coronal sections were washed with TBS and incubated for 1 h in 0.3% Triton X-100 in TBS (TBST) and 3% normal goat serum (NGS). They were then incubated with primary monoclonal mouse anti-TH antibody (1:2500, Millipore catalogue no. MAB5280, RRID:AB_2201526) diluted in TBST containing 1% NGS overnight (4 °C). Then, slices were incubated with a green fluorescent conjugated goat anti-mouse Alexa 488 antibody (1:500, Invitrogen catalogue no. A-11029, RRID:AB_138404) for 1h30 at room temperature. They were finally mounted on superfrost glass slides (Thermo Scientific, Illkirch, France), with Aqua-Poly/Mount (Polysciences Inc., Hirschberg an der Bergstraße, Germany). Fluorescent acquisitions of TH labelling and mCherry expression were taken with an objective x20 / NA 0.8 on a slide scanner (Axioscan Z1, Zeiss, Göttingen, Germany), and analyzed with ImageJ. Fluorescent illustrations were taken with an objective x20 / NA 0.8 on a spinning-disk confocal microscope (CSU-W1 confocal scanner unit Yokogawa (Gataca Systems, Massy, France), Prime 95B sCMOS Camera (Teledyne Photometrics, Birmingham, UK) and microscope Axio-observer Z1(Zeiss)). Z-stacks of digital images were captured using ZEN software (Zeiss).

### Statistical analyses

Parametric analyses were performed with assumptions of normality (Shapiro-Wilk and Kolmogorov-Smirnov tests). For sphericity, tests were corrected using the Greenhouse-Geisser correction (6). Data were analyzed by unpaired t-test, paired t-test, simple linear regression, one-way ANOVA, two-way ANOVAs and RM two-way ANOVAs, three-way ANOVAs, RM three-way ANOVAs depending on the experimental design, using Prism 8 (GraphPad Prism). Due to technical reasons, such as leakage or artefacts obtained in HPLC, some values were missing in the microdialysis experiment (< 2% of the value obtained in this experiment). In this case, data were analyzed by fitting a mixed model proposed by the statistical software. This mixed model uses a compound symmetry covariance matrix and is fit using Restricted Maximum Likelihood (REML). When indicated, *post hoc* analyses were carried out with the Bonferroni’s correction procedure. Significance for p values was set at α = 0.05. Effect sizes for the ANOVAs were also reported using partial η^2^ values (1,7). Determining these values from the mixed-model analysis was however not accessible.

## SUPPLEMENTAL FIGURES

**Supplementary Fig. S1.**
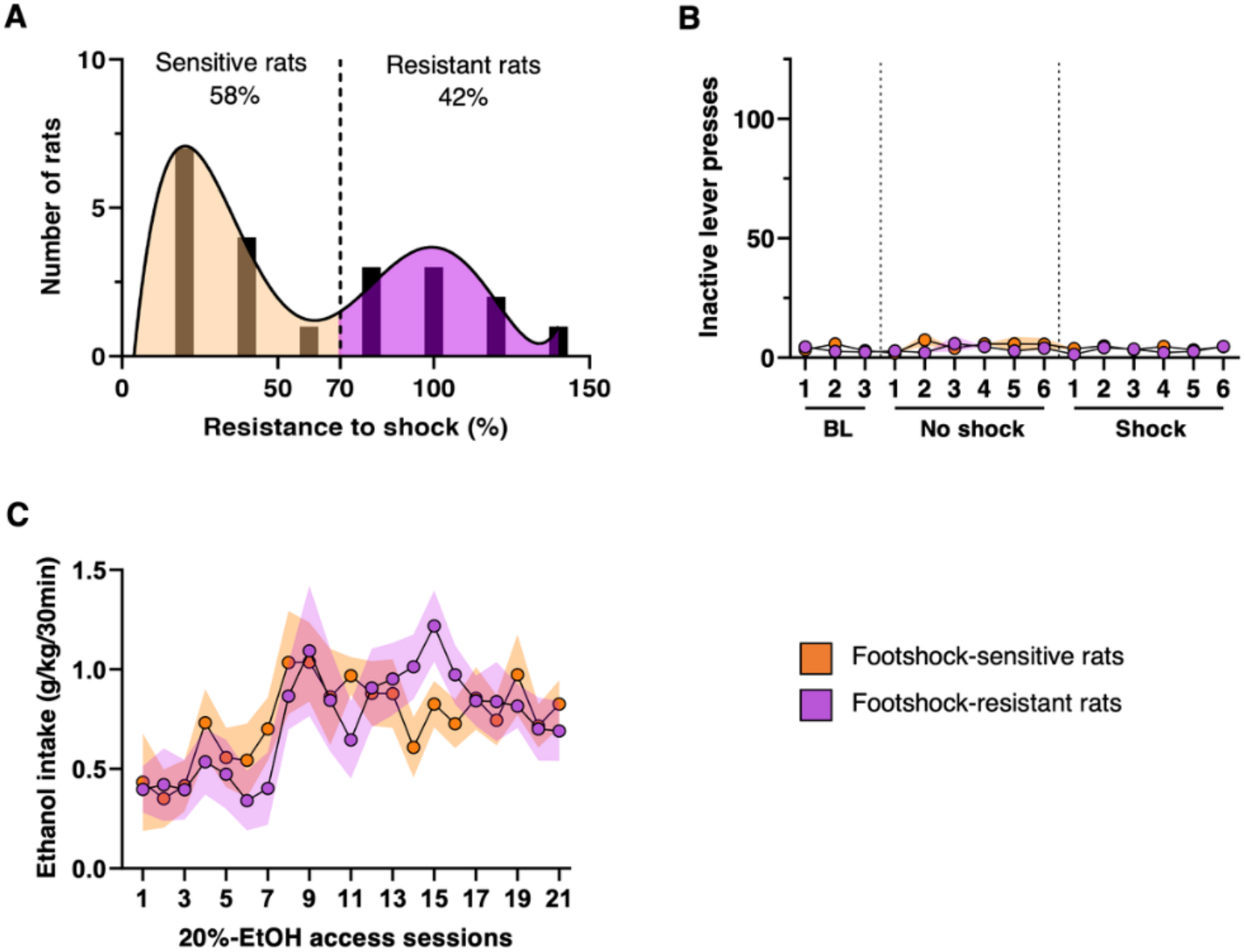
Behavioral characterization of footshock-sensitive and footshock-resistant rats. **(A)** Bimodal distribution of the population of rats self-administering alcohol under footshock-punishment sessions. 42% of rats were *footshock-resistant* (*n*=9) and 58% were *footshock-sensitive* (*n*=12). **(B)** Number of inactive lever presses in 30-min self-administration sessions of 20% EtOH (FR3), during baseline, “no-shock” and “shock” sessions. RM two-way ANOVAs found no effect of treatment, transgene or treatment × transgene interaction [F_s_<1.06, P>0.39, partial η^2^<0.05]. **(C)** Ethanol intake during the first 30 minutes of intermittent-access 20%-ethanol two-bottle-choice sessions (*IA 20%-EtOH 2BC*). RM two-way ANOVAs showed a significant effect of session [F_(8, 158)_=3.41, P<0.001, partial η^2^=0.15) but neither effect of group nor session × group interaction [F_s_<0.72, P>0.5, partial η^2^<0.04]. Data are expressed in mean ± SEM.

**Supplementary Fig. S2.**
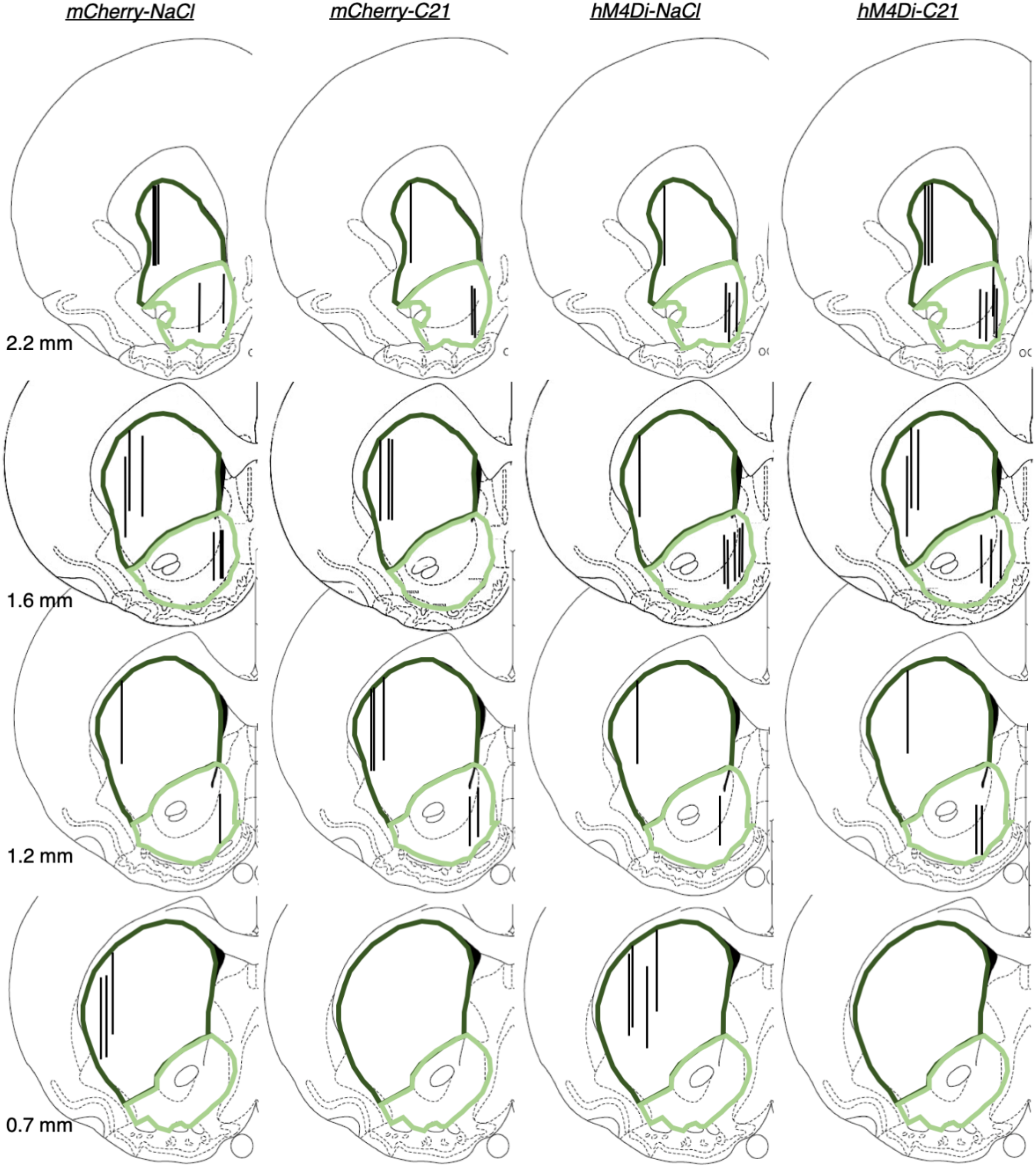
Schematic representation of the microdialysis probe placement across groups. The location of the dialysis membrane in each individual is represented by vertical bars. The dorsal striatum and the nucleus accumbens are delineated in dark and light green respectively. The numbers indicate the distance anterior to bregma (mm). Coronal sections were taken from (3).

**Supplementary Fig. S3.**
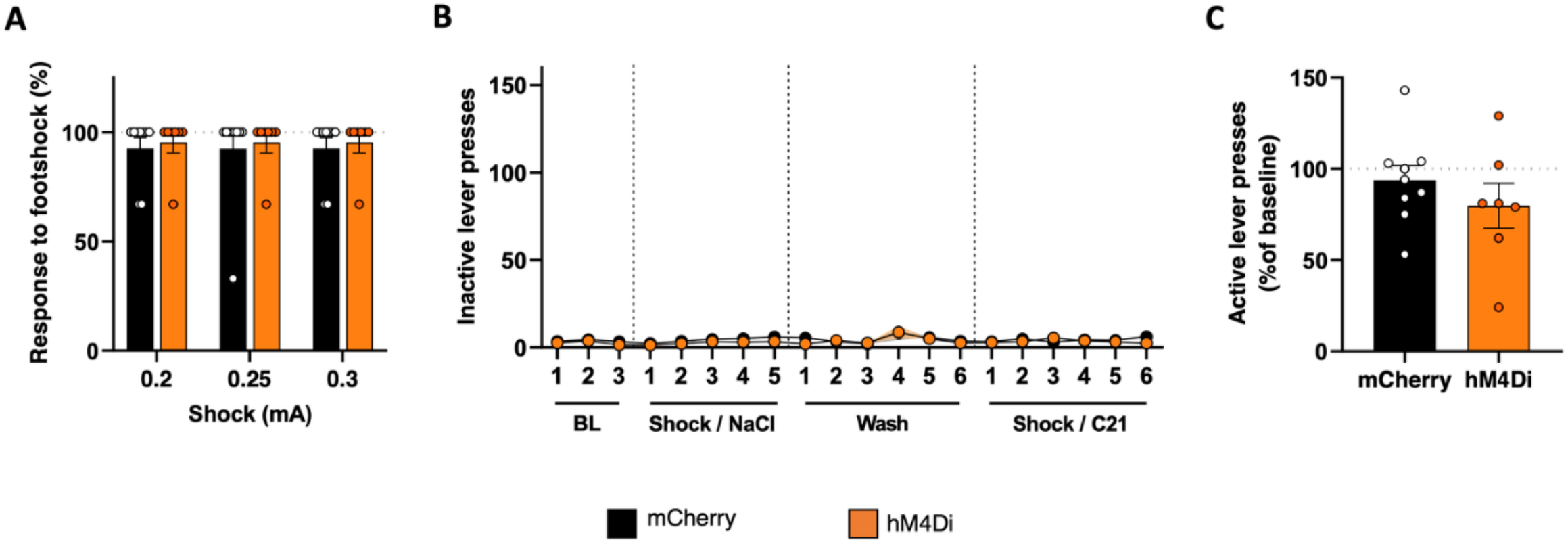
Chemogenetically-induced nigrostriatal hypodopaminergia does not decrease sensitivity to footshocks and does not increase inactive lever presses or operant ethanol self-administration. **(A)** In order to ensure that chemogenetic did not decrease sensitivity to footshock, we evaluate the effect of C21 on the percentage of response to footshocks in absence of alcohol, at intensity used during self-administration of 20%-EtOH (0.25 mA) and at two surrounding intensities (0.2 and 0.3 mA) in *hM4Di* rats (orange, *n*=7) and *mCherry* rats (black, *n*=10). RM two-way ANOVAs found no effect of the group, intensity or group intensity interaction [F_s_<0.41, P>0.5, partial η^2^<0.01]. **(B)** Number of inactive lever presses in 30-min self-administration sessions of 20%-EtOH (FR3), during baseline, “Shock/NaCl”, “Wash” and “Shock/C21” sessions. RM two-way ANOVA showed no effect of session, transgene or session × transgene interaction [F_s_<1.95, P>0.1, partial η^2^<0.12]. **(C)** Effect of C21 on the percentage of lever presses over the course of a 30-min self-administration session of 20%-EtOH (FR3) without shock normalized to baseline. Unpaired t-test showed no difference of performances between the two groups [t=0.98, P=0.34]. Data are expressed in mean ± SEM.

**Supplementary Fig. S4.**
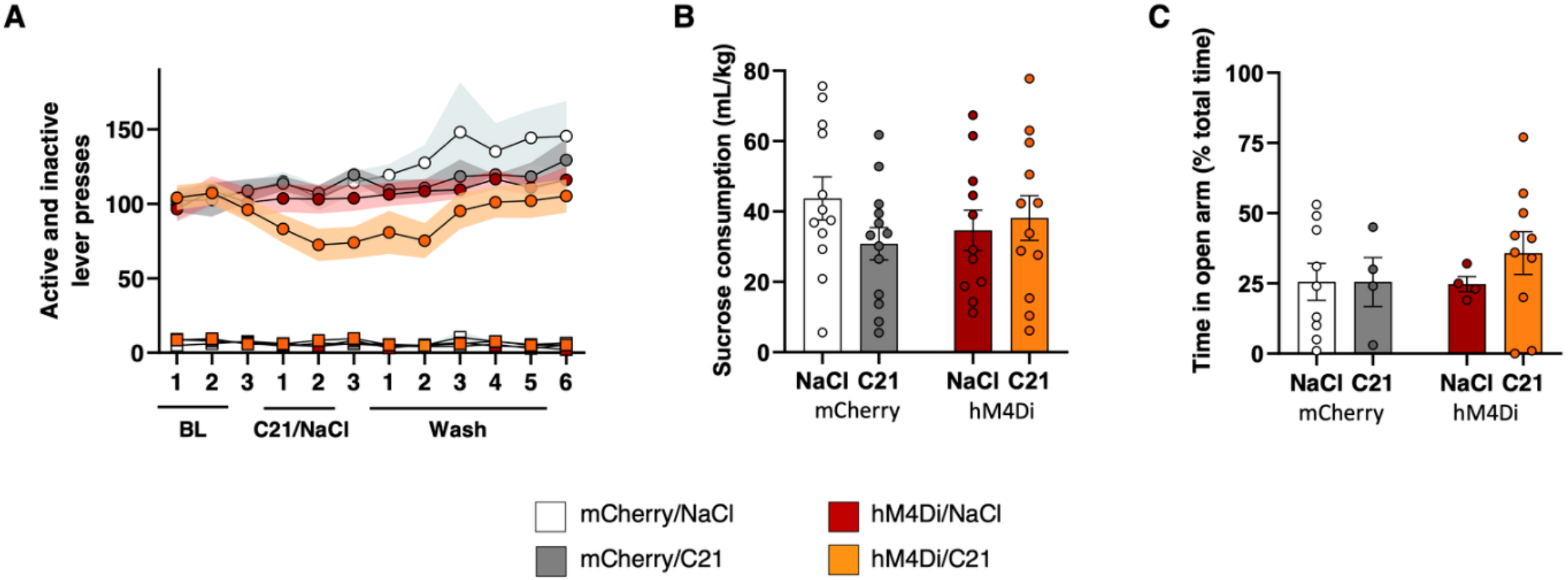
Nigrostriatal hypodopaminergia does not altered reward sensitivity or induce anxiety-related behavior. **(A)** Number of active (circle) and inactive (square) lever presses to obtained 2.5%-sucrose deliveries in 60-min self-administration sessions under continuous reinforcement (FR1), during baseline, “C21/NaCl” and “Wash” sessions, in *hM4Di* rats treated with C21 (orange, *n*=8) or NaCl (red, *n*=9) and *mCherry* rats treated with C21 (grey, *n*=10) or NaCl (white, *n*=10). On active lever presses, RM three-way ANOVAs found a significant effect of session and transgene [Fs>4.01; P<0.05; partial η^2^>0.11] but no effect of treatment or any interaction between these factors [Fs<2.42; P>0.1; partial η^2^<0.06]. On inactive lever presses, RM three-way ANOVAs revealed a significant effect of session [F_(7, 218)_=2.65; P<0.05; partial η^2^<0.07] but no effect of treatment, transgene, or any interaction between these factors [Fs<1.14; P>0.33; partial η^2^<0.33]. **(B)** Sucrose consumption during a 60-min 2.5%-sucrose two-bottle-choice drinking session by *hM4Di* rats treated with C21 (orange, *n*=12) or NaCl (red, *n*=11) and *mCherry* rats treated with C21 (grey, *n*=13) or NaCl (white, *n*=12). **(C)** Percentage of time spent in open arms of an elevated-plus maze by *hM4Di* rats treated with C21 (orange, *n*=10) or NaCl (red, *n*=4) and *mCherry* rats treated with C21 (grey, *n*=4) or NaCl (white, *n*=9). Two-way ANOVAs showed no effect of treatment, transgene or treatment × transgene interaction in these two tests [F_s_<2.04, P>0.16, partial η^2^<0.04]. Data are expressed in mean ± SEM.

**Table S1. Summary of statistical analyses**

## Notes

### Competing Interest Statement

The authors have declared no competing interest.

### Summary of Updates

This version of the manuscript has been revisited to update discussion and to include in the manuscript a part of the material and methods.

