## Supplemental Table S1 for "Hypodopaminergic state of the nigrostriatal pathway drives compulsive alcohol use"

| Figures | Questions ? | Answ. | Tests | Normality | Results | a | partial $\eta^2$ | Post-hoc | Location |
|---|---|---|---|---|---|---|---|---|---|
| Fig. 1B | Is there an effect of session? | Yes | RM Two-way ANOVA | No | F(6, 109) = 7.82 | <0.001 | 0,29 | | |
| | Is there an effect of group? | No | | | F(1, 19) = 0.13 | 0.72 | 0,01 | | |
| | Is there a session x group interaction? | No | | | F(20, 380) = 0.47 | 0.98 | 0,02 | | |
| Fig. 1C | Is there an effect of session? | Yes | RM Two-way ANOVA | No | F(4, 81) = 5.345 | <0.001 | 0,22 | Bonferonni | Shock session 6 ** |
| | Is there an effect of group? | No | | | F(1, 19) = 1.63 | 0.22 | 0,05 | | |
| | Is there a session x group interaction? | Yes | | | F(14, 266) = 2.86 | <0.001 | 0,13 | | |
| Fig. 1D | Is there an effect of session? | Yes | RM Two-way ANOVA | Yes | F(1, 19) = 44.67 | <0.001 | 0,61 | Bonferonni | Shock *** |
| | Is there an effect of group? | Yes | | | F(1, 19) = 25.25 | <0.001 | 0,70 | | |
| | Is there a session x group interaction? | Yes | | | F(1, 19) = 20.89 | <0.001 | 0,52 | | |
| Fig. 1E DLS | Is there an effect of group? | Yes | One-way ANOVA | Yes | F(2, 26) = 5.56 | <0.01 | 0,30 | Bonferonni | S vs R ** |
| Fig. 1E DMS | Is there an effect of group? | No | One-way ANOVA | Yes | F(2, 25) = 0.55 | 0.58 | 0,04 | | |
| Fig. 1E NAc | Is there an effect of group? | No | One-way ANOVA | Yes | F(2, 26) = 0.26 | 0.77 | 0,02 | | |
| Fig. 1F DLS | Is there a correlation between the resistance to footshock and the DA level of the DLS? | Yes | Simple linear regression | | F(1, 19) = 16.18 | <0.001 | | | |
| Fig. 1F DMS | Is there a correlation between the resistance to footshock and the DA level of the DMS? | No | Simple linear regression | | F(1, 18) = 1.14 | 0.3 | | | |
| Fig. 1F NAc | Is there a correlation between the resistance to footshock and the DA level of the NAc? | No | Simple linear regression | | F(1, 19) = 0.04 | 0.85 | | | |
| Fig. 2C | Is there a difference in the viral expression depending of area? | Yes | Three-way ANOVA | Yes | F(2, 129) = 158 | < 0.001 | 0,71 | | |
| | Is there a difference in the viral expression depending of transgene? | No | | | F(1, 129) = 0.54 | 0.46 | 0,004 | | |
| | Is there a difference in the viral expression depending of treatment? | No | | | F(1, 129) = 0.01 | 0.91 | 0,0001 | | |
| | Is there a area x transgene interaction? | No | | | F(2, 129) = 1.04 | 0.36 | 0,02 | | |
| | Is there a area x treatment interaction? | No | | | F(2, 129) = 0.37 | 0.69 | 0,006 | | |
| | Is there a transgene x treatment interaction? | No | | | F(1, 129) = 0.15 | 0.7 | 0,001 | | |
| | Is there a area x transgene x treatment interaction? | No | | | F(2, 129) = 0.08 | 0.93 | 0,001 | | |
| Fig. 2E | Is there an effect of treatment? | No | Mixed-effects model (REML) | No | F(1, 27) = 0.59 | 0.45 | n.a. | | |
| | Is there an effect of transgene? | No | | | F(1, 27) = 0.54 | 0.47 | n.a. | | |
| | Is there an effect of time? | No | | | F(2, 65) = 1.76 | 0.17 | n.a. | | |
| | Is there a treatment x transgene interaction? | No | | | F(1, 27) = 0.04 | 0.85 | n.a. | | |
| | Is there a treatment x time interaction? | No | | | F(7, 185) = 0.73 | 0.65 | n.a. | | |
| | Is there a transgene x time interaction? | No | | | F(7, 185) = 0.96 | 0.46 | n.a. | | |
| | Is there a treatment x transgene x time interaction? | No | | | F(7, 185) = 1.237 | 0.28 | n.a. | | |
| Fig. 2F | Is there an effect of treatment? | No | Mixed-effects model (REML) | No | F(1, 25) = 0.04 | 0.85 | n.a. | | |
| | Is there an effect of transgene? | No | | | F(1, 25) = 0.15 | 0.70 | n.a. | | |
| | Is there an effect of time? | No | | | F(4, 103) = 1.3 | 0.27 | n.a. | | |
| | Is there a treatment x transgene interaction? | No | | | F(1, 25) = 0.08 | 0.79 | n.a. | | |
| | Is there a treatment x time interaction? | No | | | F(7, 171) = 0.18 | 0.99 | n.a. | | |
| | Is there a transgene x time interaction? | No | | | F(7, 171) = 0.35 | 0.93 | n.a. | | |
| | Is there a treatment x transgene x time interaction? | No | | | F(7, 171) = 1.02 | 0.42 | n.a. | | |
| Fig. 2G | Is there an effect of treatment? | Yes | Two-way ANOVA | Yes | F(1, 27) = 8.71 | <0.01 | 0,24 | Bonferonni | hM4Di-C21 vs all : ** |
| | Is there an effect of transgene? | Yes | | | F(1, 27) = 8.26 | <0.01 | 0,23 | | |
| | Is there a treatment x transgene interaction? | Yes | | | F(1, 27) = 8.63 | <0.01 | 0,24 | | |

| | | | | | | | | | |
|---|---|---|---|---|---|---|---|---|---|
| Fig. 2H | Is there an effect of treatment? | No | Two-way ANOVA | No | F(1, 25) = 0.29 | 0.59 | 0,01 | | |
| | Is there an effect of transgene? | No | | | F(1, 25) = 0.11 | 0.74 | 0,001 | | |
| | Is there a treatment x transgene interaction? | No | | | F(1, 25) = 0.03 | 0.87 | 0,001 | | |
| Fig. 3B | Is there a difference in the viral expression depending of area? | Yes | Two-way ANOVA | No | F(2, 90) = 116.4 | < 0.001 | 0,72 | | |
| | Is there a difference in the viral expression depending of transgene? | No | | | F(1, 90) = 0.87 | 0.35 | 0,01 | | |
| | Is there a area x transgene interaction? | No | | | F(2, 90) = 1.99 | 0.14 | 0,04 | | |
| Fig. 3C | Is there an effect of session? | Yes | RM Two-way ANOVA | No | F(4, 59) = 10.56 | <0.001 | 0,43 | Bonferroni | No |
| | Is there an effect of transgene? | No | | | F(1, 14) = 0.1 | 0.76 | 0,01 | | |
| | Is there a session x transgene interaction? | No | | | F(23, 308) = 0.57 | 0.94 | 0,04 | | |
| Fig. 3D | Is there an effect of session? | Yes | RM Two-way ANOVA | No | F(4, 59) = 21.55 | <0.001 | 0,61 | | |
| | Is there an effect of transgene? | No | | | F(1, 14) = 0.02 | 0.9 | 0,001 | | |
| | Is there a session x transgene interaction? | Yes | | | F(19, 266) = 2.18 | < 0.01 | 0,13 | | |
| Fig. 3E | Is there an effect of session? | Yes | RM Two-way ANOVA | Yes | F(1, 14) = 9.69 | <0.01 | 0,41 | Bonferroni | mCherry vs hM4Di during C21 and NaCl vs C21 in hM4Di = *** |
| | Is there an effect of transgene? | Yes | | | F(1, 14) = 15 | <0.01 | 0,83 | | |
| | Is there a session x transgene interaction? | Yes | | | F(1, 14) = 18.41 | <0.001 | 0,57 | | |
| Fig. 3F | Is there a difference between Shock/NaCl an Shock/C21 periods in hM4Di-expressing rats? | Yes | Paired t-test | Yes | t = 4.3, df = 6 | <0.01 | | | |
| | Is there a difference between Shock/NaCl an Shock/C21 periods in mCherry-expressing rats? | No | Paired t-test | Yes | t = 1.02, df = 8 | 0.34 | | | |
| Fig. 3G | Is there a correlation between difference of lever presses during Shock/C21 and Shock/NaCl periods and hM4Di expression in the dSNc? | Yes | Simple linear regression | | F(1, 5) = 7.64 | <0.05 | | | |
| Fig. 3H | Is there a correlation between difference of lever presses during Shock/C21 and Shock/NaCl periods and hM4Di expression in the mSNc? | No | Simple linear regression | | F(1, 5) = 0.18 | 0.69 | | | |
| Fig. 4B | Is there a difference in the viral expression depending of area? | Yes | Three-way ANOVA | Yes | F(2, 140) = 346.7 | < 0.01 | 0,83 | | |
| | Is there a difference in the viral expression depending of transgene? | No | | | F(1, 70) = 0.35 | 0.55 | 0,01 | | |
| | Is there a difference in the viral expression depending of treatment? | No | | | F(1, 70) = 0.16 | 0.69 | 0,003 | | |
| | Is there a area x transgene interaction? | No | | | F(2, 140) = 1.31 | 0.27 | 0,02 | | |
| | Is there a area x treatment interaction? | No | | | F(2, 140) = 0.05 | 0.95 | < 0.001 | | |
| | Is there a transgene x treatment interaction? | No | | | F(1, 70) = 1.13 | 0.29 | 0,02 | | |
| | Is there a area x transgene x treatment interaction? | No | | | F(2, 140) = 1.11 | 0.33 | 0,02 | | |
| Fig. 4C | Is there an effect of treatment? | No | RM-Three-way ANOVA | No | F(1, 33) = 1.84 | 0.18 | 0,10 | Bonferroni | No |
| | Is there an effect of transgene? | No | | | F(1, 33) = 2.76 | 0.11 | 0,15 | | |
| | Is there an effect of session? | Yes | | | F(4, 145) = 6.71 | < 0.001 | 0,17 | | |
| | Is there a treatment x transgene interaction? | No | | | F(1, 33) = 2.69 | 0.11 | 0,14 | | |
| | Is there a treatment x session interaction? | Yes | | | F(11, 363) = 3.41 | < 0.001 | 0,09 | | |
| | Is there a transgene x session interaction? | Yes | | | F(11, 363) = 2.96 | <0.001 | 0,1 | | |
| | Is there a treatment x transgene x session interaction? | Yes | | | F(11,363) = 2.67 | <0.01 | 0,07 | | |
| Fig. 4D | Is there an effect of treatment? | Yes | Two-way ANOVA | Yes | F(1, 33) = 10.79 | < 0.01 | 0,25 | Bonferroni | hM4Di-C21 vs all (***) |
| | Is there an effect of transgene? | Yes | | | F(1, 33) = 17.95 | < 0.001 | 0,35 | | |
| | Is there a treatment x transgene interaction? | Yes | | | F(1, 33) = 13.38 | < 0.001 | 0,29 | | |

| | | | | | | | |
|---|---|---|---|---|---|---|---|
| Fig. 4E | Is there an effect of treatment? | No | RM-Three-way ANOVA | Yes | F(1, 44) = 0.23 | 0.63 | 0,01 |
| | Is there an effect of transgene? | No | | | F(1, 44) = 0.92 | 0.34 | 0,04 |
| | Is there an effect of time? | Yes | | | F(4, 175) = 380.4 | < 0.001 | 0,90 |
| | Is there a treatment x transgene interaction? | No | | | F(1, 44) = 0.27 | 0.61 | 0,01 |
| | Is there a treatment x time interaction? | Marginal | | | F(5, 220) = 2.24 | 0.05 | 0,05 |
| | Is there a transgene x time interaction? | No | | | F(5, 220) = 0.44 | 0.82 | 0,01 |
| | Is there a treatment x transgene x time interaction? | No | | | F(5, 220) = 1.3 | 0.26 | 0,03 |
| Fig. 4F | Is there an effect of treatment? | No | RM-Three-way ANOVA | Yes | F(1, 44) = 1.51 | 0.23 | 0,07 |
| | Is there an effect of transgene? | No | | | F(1, 44) = 0.45 | 0.51 | 0,02 |
| | is there an effect of side? | Yes | | | F(1, 44) = 10.27 | < 0.01 | 0,19 |
| | Is there a treatment x transgene interaction? | No | | | F(1, 44) = 0.14 | 0.72 | 0,01 |
| | Is there a treatment x side interaction? | No | | | F(1, 44) = 0.0002 | 0.99 | < 0.001 |
| | Is there a transgene x side interaction? | No | | | F(1, 44) = 0.29 | 0.59 | 0,01 |
| | Is there a treatment x transgene x side interaction? | Marginal | | | F(1, 44) = 3.55 | 0.07 | 0,07 |
| Fig. 4G | Is there an effect of treatment? | No | Two-way ANOVA | No | F(1, 44) = 1.7 | 0.2 | 0,04 |
| | Is there an effect of transgene? | No | | | F(1, 44) = 0.36 | 0.55 | 0,01 |
| | Is there a treatment x transgene interaction? | No | | | F(1, 44) = 0.13 | 0.72 | 0,003 |
| Fig. 4H | Is there an effect of treatment? | No | Two-way ANOVA | Yes | F(1, 44) = 0.0003 | 0.99 | < 0.001 |
| | Is there an effect of transgene? | No | | | F(1, 44) = 1.49 | 0.23 | 0,03 |
| | Is there a treatment x transgene interaction? | Marginal | | | F(1, 44) = 3.68 | 0.06 | 0,08 |
| Fig. 4I | Is there an effect of treatment? | No | Two-way ANOVA | Yes | F(1, 44) = 0.6 | 0.44 | 0,01 |
| | Is there an effect of transgene? | No | | | F(1, 44) = 1.85 | 0.18 | 0,04 |
| | Is there a treatment x transgene interaction? | No | | | F(1, 44) = 0.29 | 0.6 | 0,01 |
| fig. S1B | Is there an effect of session? | No | RM Two-way ANOVA | No | F(5, 89) = 1.04 | 0.4 | 0,05 |
| | Is there an effect of trait? | No | | | F(1, 19) = 0.50 | 0.49 | 0,0204 |
| | Is there a session x trait interaction? | No | | | F(14, 266) = 1,06 | 0.39 | 0,05 |
| fig. S1C | Is there an effect of session? | Yes | RM Two-way ANOVA | No | F(8, 158) = 3.41 | <0.001 | 0,15 |
| | Is there an effect of trait? | No | | | F(1, 19) = 0.02 | 0.9 | 0,0002 |
| | Is there a session x trait interaction? | No | | | F(20, 380) = 0.72 | 0.81 | 0,04 |
| fig. S3A | Is there an effect of intensity? | No | RM Two-way ANOVA | No | F(2, 24) < 0.001 | > 0.99 | <0.001 |
| | Is there an effect of transgene? | No | | | F(1, 14) = 0.41 | 0.53 | 0,01 |
| | Is there a intensity x transgene interaction? | No | | | F(2, 28) <0.001 | > 0.99 | <0.001 |
| fig. S3B - Inactive lever presses | Is there an effect of session? | No | RM Two-way ANOVA | No | F(5, 64) =1.95 | 0.1 | 0,12 |
| | Is there an effect of transgene? | No | | | F(1, 14) = 0.82 | 0.38 | 0,02 |
| | Is there a session x transgene interaction? | No | | | F(19, 266) = 0.53 | 0.949 | 0,036 |
| fig. S3C | Is there an effect of treatment? | No | Unpaired t-test | Yes | t = 0.98, df = 14 | 0.34 | |
| fig. S4A - Active lever Presses | Is there an effect of treatment? | No | RM-Three-way ANOVA | No | F(1, 33) = 2.42 | 0.13 | 0,06 |
| | Is there an effect of transgene? | Yes | | | F(1, 33) = 5.49 | < 0.05 | 0,13 |
| | Is there an effect of session? | Yes | | | F(2, 64) = 4.01 | < 0.05 | 0,11 |
| | Is there a treatment x transgene interaction? | No | | | F(1, 33) = 0.18 | 0.68 | 0,005 |
| | Is there a treatment x session interaction? | No | | | F(11, 363) = 0.99 | 0.45 | 0,03 |
| | Is there a transgene x session interaction? | No | | | F(11, 363) = 1.58 | 0.10 | 0,05 |
| | Is there a treatment x transgene x session interaction? | No | | | F(11,363) = 0.99 | 0.46 | 0,03 |

| | | | | | | | |
|---|---|---|---|---|---|---|---|
| fig. S4A - Inactive lever Presses | Is there an effect of treatment? | No | RM-Three-way ANOVA | No | F(1, 33) = 0.02 | 0.88 | 0,001 |
| | Is there an effect of transgene? | No | | | F(1, 33) = 0.03 | 0.86 | 0,001 |
| | Is there an effect of session? | Yes | | | F(7, 218) = 2.65 | < 0.05 | 0,07 |
| | Is there a treatment x transgene interaction? | No | | | F(1, 33) = 0.58 | 0.45 | 0,013 |
| | Is there a treatment x session interaction? | No | | | F(11, 363) = 1.14 | 0.33 | 0,033 |
| | Is there a transgene x session interaction? | No | | | F(11, 363) = 0.81 | 0.63 | 0,024 |
| | Is there a treatment x transgene x session interaction? | No | | | F(11,363) = 0.76 | 0.68 | 0,023 |
| fig. S4B | Is there an effect of treatment? | No | Two-way ANOVA | Yes | F(1, 44) = 0.66 | 0.42 | 0,01 |
| | Is there an effect of transgene? | No | | | F(1, 44) = 0.02 | 0.88 | < 0.001 |
| | Is there a treatment x transgene interaction ? | No | | | F(1, 44) = 2.04 | 0.16 | 0,04 |
| fig. S4C | Is there an effect of treatment? | No | Two-way ANOVA | No | F(1, 23) = 0.42 | 0.52 | 0,02 |
| | Is there an effect of transgene? | No | | | F(1, 23) = 0.31 | 0.58 | 0,01 |
| | Is there a treatment x transgene interaction? | No | | | F(1, 23) = 0.43 | 0.52 | 0,02 |
